# A Mouse Model of the Protease Activated Receptor 4 (PAR4) Pro310Leu Variant has Reduced Platelet Reactivity

**DOI:** 10.1101/2023.12.01.569075

**Authors:** Xu Han, Elizabeth A. Knauss, Maria de la Fuente, Wei Li, Ronald A Conlon, David F. LePage, Weihong Jiang, Stephanie A. Renna, Steven E. McKenzie, Marvin T. Nieman

**Author notes:** Denotes equal contribution (Xu Han, Elizabeth A. Knauss). Corresponding author Marvin T. Nieman, Department of Pharmacology, Case Western Reserve University, 2109 Adelbert Road W309B, Cleveland, OH, 44106-4965, USA.

## Abstract

**Background:** Protease activated receptor 4 (PAR4) mediates thrombin signaling on platelets and other cells. Our recent structural studies demonstrated a single nucleotide polymorphism in extracellular loop 3 (ECL3), PAR4-P310L (rs2227376) leads to a hypo-reactive receptor.

**Objectives:** The goal of this study was to determine how the hypo-reactive PAR4 variant in ECL3 impacts platelet function in vivo using a novel knock-in mouse model (PAR4-322L).

**Methods:** A point mutation was introduced into the PAR4 gene, F2rl3, via CRISPR/Cas9 to create PAR4-P322L, the mouse homolog to human PAR4-P310L. Platelet response to PAR4 activation peptide (AYPGKF), thrombin, ADP, and convulxin was monitored by αIIbβ3 integrin activation and P-selectin translocation using flow cytometry or platelet aggregation. In vivo responses were determined by the tail bleeding assay and the ferric chloride-induced carotid artery injury model.

**Results:** PAR4-P/L and PAR4-L/L platelets had a reduced response to AYPGKF and thrombin measured by P-selectin translocation or αIIbβ3 activation. The response to ADP and convulxin was unchanged among genotypes. In addition, both PAR4-P/L and PAR4-L/L platelets showed a reduced response to thrombin in aggregation studies. There was an increase in the tail bleeding time for PAR4-L/L mice. The PAR4-P/L and PAR4-L/L mice both showed an extended time to arterial thrombosis.

**Conclusions:** PAR4-322L significantly reduced platelet responsiveness to AYPGKF and thrombin, which is in agreement with our previous structural and cell signaling studies. In addition, PAR4-322L had prolonged arterial thrombosis time. Our mouse model provides a foundation to further evaluate the role of PAR4 in other pathophysiological contexts.

**Essentials:**

- A mouse model was created to represent the PAR4-P310L sequence variant.
- PAR4-P322L leads to reduced platelet reactivity in response to PAR4-activation peptide and thrombin, while the ADP and GPVI signaling pathways were unaffected.
- The PAR4-P322L mutation decreases time to occlusion in a mouse model of arterial thrombosis.
- The PAR4-P322L mouse model provides a foundation to further explore the role of PAR4 in hemostasis and thrombosis.

## Introduction

Platelets play a pivotal role in primary hemostasis, thrombosis, inflammation, and vascular biology. These anuclear discoid cells circulate in the bloodstream to patrol the integrity of the vascular system.[1] Upon injury, platelets quickly activate, change shape, release granule contents, and aggregate to form the hemostatic plug in the presence of fibrinogen. Platelet activation can be triggered by many physiological agonists, including thrombin, the most potent platelet agonist and a key protease in coagulation.[2,3] Thrombin signals through two protease activated receptors (PARs), PAR1 and PAR4, on the surface of human platelets.[4] PARs belong to the GPCR superfamily and have a unique activation mechanism whereby the N-terminus is enzymatically cleaved to unmask the tethered ligand.[5] The new N-terminus interacts with the endogenous ligand binding site to induce a global structural rearrangement that activates downstream signaling.[6,7] PAR1 and PAR4 both signal through Gα_q_ and Gα_12/13_, however with different kinetics.[8] PAR1 leads to rapid signaling that is quickly dissipated, whereas PAR4 leads to prolonged signaling.[4,9,10] This sustained signaling associated with PAR4 activation is essential for thrombosis, highlighting PAR4 as a promising target for antiplatelet therapies.[11–13]

Over the past 10 years, platelet thrombin receptors have been appealing targets for antiplatelet therapies, which led to the first-in-class FDA approved PAR1 inhibitor, vorapaxar. However, targeting PAR1 comes with a significant risk of bleeding, which outweighs its clinical benefits in preventing cardiovascular events.[14,15] In recent years, PAR4 has become a rising star as a safer antiplatelet and antithrombotic target for a number of reasons. First, targeting PAR4 signaling without inhibiting PAR1 allows platelets to continue to respond to low levels of thrombin and preserves normal hemostasis.[12] Second, pharmacological inhibition of PAR4 prevents thrombin-mediated PAR4 activation at high concentrations that are associated with pathological thrombosis. Third, since the prolonged signaling mediated by PAR4 activation is associated with factor V release from α-granules and microparticle generation[16], selectively inhibiting PAR4 would not only prevent thrombus formation but also reduce platelet procoagulant activity.[11] Collectively, this has culminated in the development of a number of PAR4 antagonists in the form of pepducins, small molecule compounds, and function-blocking antibodies.[17] Two small molecule PAR4 inhibitors from Bristol Myers Squibb, BMS-986120 and BMS-986141, were the subject of clinical trials and proved to be efficient in preventing cardiovascular events with a good safety profile.[12,13,18] PAR4’s unique properties have made it an attractive therapeutic target to prevent thrombosis without hindering normal hemostasis.

Therefore, it is essential that we fully understand the mechanisms underlying PAR4 signaling. The tethered ligand mechanism was proposed in 1991 by Coughlin and colleagues, however the molecular mechanism of receptor activation is only recently understood. Recently, we used amide hydrogen/deuterium exchange (H/D exchange) mass spectrometry (MS) with purified full-length PAR4 to examine the conformational dynamics of the tethered ligand mechanism following activation by thrombin.[19] This study revealed that PAR4 activation requires a coordinated rearrangement of extracellular loop 3 (ECL3) and threonine at position 153 in the ligand binding site formed by transmembrane domain 3 (TM3) and TM7. Within ECL3, there is a single nucleotide polymorphism (SNP) in which the proline at 310 is replaced with a leucine (PAR4-310P/L, rs2227376). This natural sequence variant of PAR4 had significantly lower receptor reactivity, as measured by calcium mobilization in HEK293 cells. Natural sequence variants of PAR4 (e.g. rs773902 (PAR4-120A/T), and rs2227346 (PAR4-296F/V)) affect the receptor reactivity and subsequent platelet function.[20–23] Therefore, we hypothesize the hypo-reactive PAR4-P310L variant would reduce platelet responsiveness to thrombin stimulation

To test the impact of the PAR4-310P/L polymorphism in vivo, we used CRISPR/Cas9 to introduce a point mutation into PAR4 to generate the mouse homolog of this variant, PAR4-P322L. PAR4-P322L significantly reduced platelet responsiveness to PAR4-activation peptide (AYPGKF) and thrombin, while ADP and GPVI signaling were not affected. Further, platelet aggregation was dramatically decreased in the platelets from mice that carried one (PAR4^P/L^, heterozygous) or two (PAR4^L/L^, homozygous) alleles of PAR4-P322L. Additionally, PAR4^L/L^ mice displayed slightly extended tail bleeding compared to wild-types. PAR4-P322L also delayed arterial occlusion in the ferric chloride-induced carotid artery thrombosis model.

## Material and Methods

### Materials

Recombinant Cas9 Nuclease and single guide RNA (sgRNA) were purchased from PNA Bio (Thousand Oaks, CA). Targeting oligonucleotide was obtained from IDT (Coralville, IA). Human α-thrombin (catalog # HCT-0020, specific activity greater than 2989 U/mg) was purchased from Haematological Technologies (Essex Junction, VT). PAR4 activation peptide, AYPGKF-NH_2_, was purchased from Tocris Bioscience (Minneapolis, MN). FITC conjugated P-selectin antibody and phycoerythrin-conjugated JON/A antibody were purchased from Emfret Analytics (Würzburg, Germany). All other reagents were from Thermo Fisher Scientific (Pittsburgh, PA) except where noted.

### Animals

All animal experiments were performed in accordance with the approval from the Case Western Reserve University Animal Ethics Committee. The PAR4-P322L mutation was introduced to the mouse genome using the CRISPR-Cas9 genome-editing system. A guide sequence, 1106/fw (CTATTCAAACCCGAGCCCTG), was validated *in vitro* (sgRNA screening system, Clontech), and obtained from PNAbio. An oligo, F2rl3 P322L 100-mer: TTTCACACCTAGCAATGTGCTGCTGGTGCTGCACTATTCAAACCTGAGCCCTGAAGCCTGGGGCAATCTCTATGGAGCCTATGTGCCCAGCCTGGCACTC (mutations underlined), designed to mutate P322 and ablate the PAM sequence of 1106/fw with a conservative base change, was obtained from IDT as a PAGE purified Ultramer. Mixtures of 100 ng/μL Cas9 nuclease; 200 ng/μL sgRNA; 400 ng/μL oligo, were electroporated into C57BL/6J fertilized oocytes as previously described.[24] Following screening through miSeq by the CWRU Genomics Core, two founder mice were then each bred to C57BL/6J mice for further proliferation. The pedigree from each founder was recorded separately. Line 1 was the offspring from the male founder and line 2 was the offspring from the female founder. PAR4 knockout mice (*F2rl3*^-/-^) were purchased from Mutant Mouse Regional Resource Centers.

### Preparation of Murine Platelets

Blood was collected from an equal number of male and female mice. For platelet rich plasma (PRP), blood was collected in sodium citrate, centrifuged at 2300 x *g* for 10 seconds, and incubated for 10 minutes at room temperature to obtain PRP. The remaining blood sample was centrifuged again at 2300 x *g* for 10 seconds and incubated for 10 minutes at room temperature to obtain more PRP. Platelet concentrations were quantified using a Coulter Counter (Beckman Coulter). Gel filtered platelets were prepared using Sepharose 2B (Sigma). Columns were packed in H_2_O and allowed to stand overnight. Before adding the PRP, 3 volumes of H_2_O and 3 volumes of HEPES Tyrode’s buffer, pH 7.4 (10 mM HEPES, 12 mM NaHCO_3_, 130 mM NaCl, 5 mM D-glucose, 5 mM KCl, 0.4 mM NaHPO_4_, 1 mM MgCl_2_) were passed through the column. The gel-filtered platelet concentrations were quantified using a Coulter Counter.

### Detecting mouse PAR4 with Western Blot

Citrated whole blood was collected from PAR4-322 P/P, P/L, L/L and PAR4 knockout mice. Murine platelet rich plasma (PRP) was prepared from the mouse whole blood as described above. After platelets were counted using a Coulter Counter, equal amounts of platelets (1X10^8^) were transferred into a new tube and spun down at 1600 x *g*. The pelleted murine platelets were lysed by RIPA buffer (Invitrogen) with protease inhibitor cocktail (Roche) on ice for 30 minutes. The lysates were further spun down at 21300 x *g* for 20 minutes at 4°C. The supernatant was transferred to a new tube and mixed with loading dye. Proteins were separated by SDS-PAGE and transferred onto a nitrocellulose membrane. Mouse PAR4 was detected using a goat polyclonal antibody (1 μg/ml) kindly provided by Dr. Steven McKenzie from Thomas Jefferson University. The primary antibody was detected with an anti-goat 800 secondary antibody (0.1 μg/ml) (LiCor). Total protein loaded was quantified by Revert 700 Total Protein Stain (LiCor) following the commercial protocol.

### Flow Cytometry

Murine PRP were used to measure the platelet reactivity in response to PAR4-activation peptide (PAR4-AP, AYPGKF-NH_2_), ADP, or convulxin. The activation of platelets was measured by the surface expression of P-selectin using FITC-conjugated P-selectin antibody and the activation of integrin αIIbβ3 using phycoerythrin-conjugated JON/A antibody (Emfret). Specifically, PRP was diluted to 5 x 10^4^ platelets/μL with HEPES Tyrode’s buffer (pH = 7.4). 10 μL of diluted PRP containing 5 x 10^5^ platelets was incubated with 5 μL FITC-conjugated P-selectin antibody, 5 μL PE-conjugated JON/A antibody and 5 μL agonist for 20 minutes in the dark at room temperature. The platelets were then fixed with 1% formaldehyde and the fluorescence were determined using BD LSRFortessa^TM^ cell analyzer (BD Biosciences, Mississauga, ON, Canada). A negative control with no antibodies was included.

Gel filtered murine platelets were used to measure the platelet reactivity in response to thrombin by flow cytometry as described above. 10 μL of gel filtered platelets was incubated with 5 μL FITC-conjugated P-selectin antibody, 5 μL PE-conjugated JON/A antibody and 5 μL agonist for 20 minutes in the dark at room temperature. The cells were then fixed by 1% formaldehyde and the fluorescence were determined by BD LSRFortessa^TM^ cell analyzer (BD Biosciences, Mississauga, ON, Canada).

### Platelet aggregation

Gel filtered murine platelets were used to measure the platelet aggregation in response to thrombin. Platelets were diluted to the final concentration of 5 x 10^7^ platelets/mL in a 300 μL reaction volume, which contains 295 μL of gel filtered platelets and 5 μL agonist. Platelet aggregation was recorded using the CHRONO-LOG® Model 700 Whole Blood/Optical Lumi-Aggregometer paired with AGGRO/LINK^®^8 program software. (Chrono-log Corporation, PA)

### Tail bleeding time assay

Both the male and female mice at 8 weeks of age were used. The mice were anesthetized by a mix of ketamine (100 mg/kg) and xylazine (10 mg/kg) via intraperitoneal injection. The body weight of the mice was recorded prior to the assay. Animals were placed in a prone position on a heater set to 37 °C. 5 mm of tail was removed by a razor blade. The tail was then immediately immersed in a 15 mL conical tube containing 15 mL pre-warmed 0.9% saline. The tail was laid vertically to the body with the tip of the tail hanging 4 cm below the heating pad and 2 cm immersed in the saline. Each mouse was monitored for 10 minutes and all re-bleeding events were recorded. Initial bleeding time was defined as the first time observing the stop of the bleeding regardless of any re-bleeding. Total bleeding time was defined as the sum of bleeding times of all bleeding on/off cycles until a stable cessation occurred (no bleeding for 60 seconds). The experiment was terminated at 10 minutes.

### Arterial thrombosis

Ferric chloride-induced carotid artery injury was performed at Marshall University by Dr. Wei Li (approved by IACUC #1033528), as previously described.[25,26] Briefly, mice 8 to 12 weeks old were anesthetized by a mixture of ketamine (100 mg/kg) and xylazine (10 mg/kg) via intraperitoneal injection. The right jugular vein was exposed and injected with 100 μl of rhodamine 6G solution (Sigma 252433-1G, 0.5 mg/ml in saline, 0.2 μm filtered) to label platelets. The injection site was ligated with a 6-0 suture to prevent bleeding. The left common carotid artery was exposed and freed from surrounding tissues. One small piece of “U” shaped black plastic was placed under the vessel to separate the carotid artery from the background fluorescence.. Saline was applied to the surgical field to keep the carotid artery moist before transferring the mouse to the attached Gibraltar Platform of a Leica DM6 FS fluorescent microscope (Deerfield, IL, USA)). A 10x water lens was positioned above the carotid artery to record baseline flow and vessel wall conditions before injury as well as clot formation after injury. Video imaging was performed using a QImaging Retiga R1: 1.4 Megapixel Color CCD camera system with mono color mode (Teledyne Photometrics, Tucson, AZ, USA)) and StreamPix version 7.1 software (Norpix, Montreal, Canada). A small piece of filter paper (1×2mm) saturated with 7.5% FeCl_3_ was placed directly onto the carotid artery at the position with the “U” shaped plastic underneath for 1 minute. After 1 minute, the filter paper was removed, the carotid artery was rinsed with saline, and then saline was reapplied between the lens and vessel. Blood flow was monitored and recorded until full occlusion or 30 minutes after FeCl_3_ injury..

## Results

### Generation of PAR4-P322L mice

Previously, we characterized a naturally occurring variant of human PAR4, PAR4-P310L (rs2227376).[19] Changing proline to leucine at position 310 in extracellular loop 3 (ECL3) significantly reduced PAR4-mediated calcium signaling in response to both PAR4-activation peptide (PAR4-AP, AYPGKF-NH_2_) and thrombin in transfected HEK293 cells. PAR4 is highly conserved between human and mouse. Human and mouse PAR4 share 74.4% sequence identity and 82.2% sequence similarity as analyzed by the Pairwise Sequence Alignment tool, EMBOSS Needle. ECL3 spans 15 amino acid residues and has 13/15 (87%) identity, and the 310 position in human PAR4 is homologous to the 322 site in mouse PAR4. Mouse PAR4-P322L also had a reduced PAR4-mediated calcium signaling in transfected HEK293 cells (data not shown).

To determine the impact of the PAR4-P310L polymorphism in vivo, we developed a mouse model using CRISPR-Cas9 to introduce the P322L mutation into mouse PAR4. The gRNAs were designed to target exon 2 from 1189 base pairs (bp) to 1287 bp of the *F2rl3* locus (**Figure 1A**). This introduced a C > T substitution at 1232 bp, which changed the proline at 322 to a leucine (**Figure 1B and C**). The homozygous PAR4-P322L mice, PAR4^L/L^, were obtained by breeding the heterozygous PAR4^P/L^ mice. Litters were born in the expected Mendelian inheritance ratios and equally divided into males and females. The platelet counts, mean platelet volume, red cell count, and leukocyte count for the PAR4^L/L^ were all unchanged compared to their PAR4^P/P^ and PAR4^P/L^ littermates. More importantly, the P322L mutation of PAR4 on the extracellular loop 3 (ECL3) did not affect protein expression on mouse platelets, validated by western blot (**Figure 1D**).

**Figure 1.**
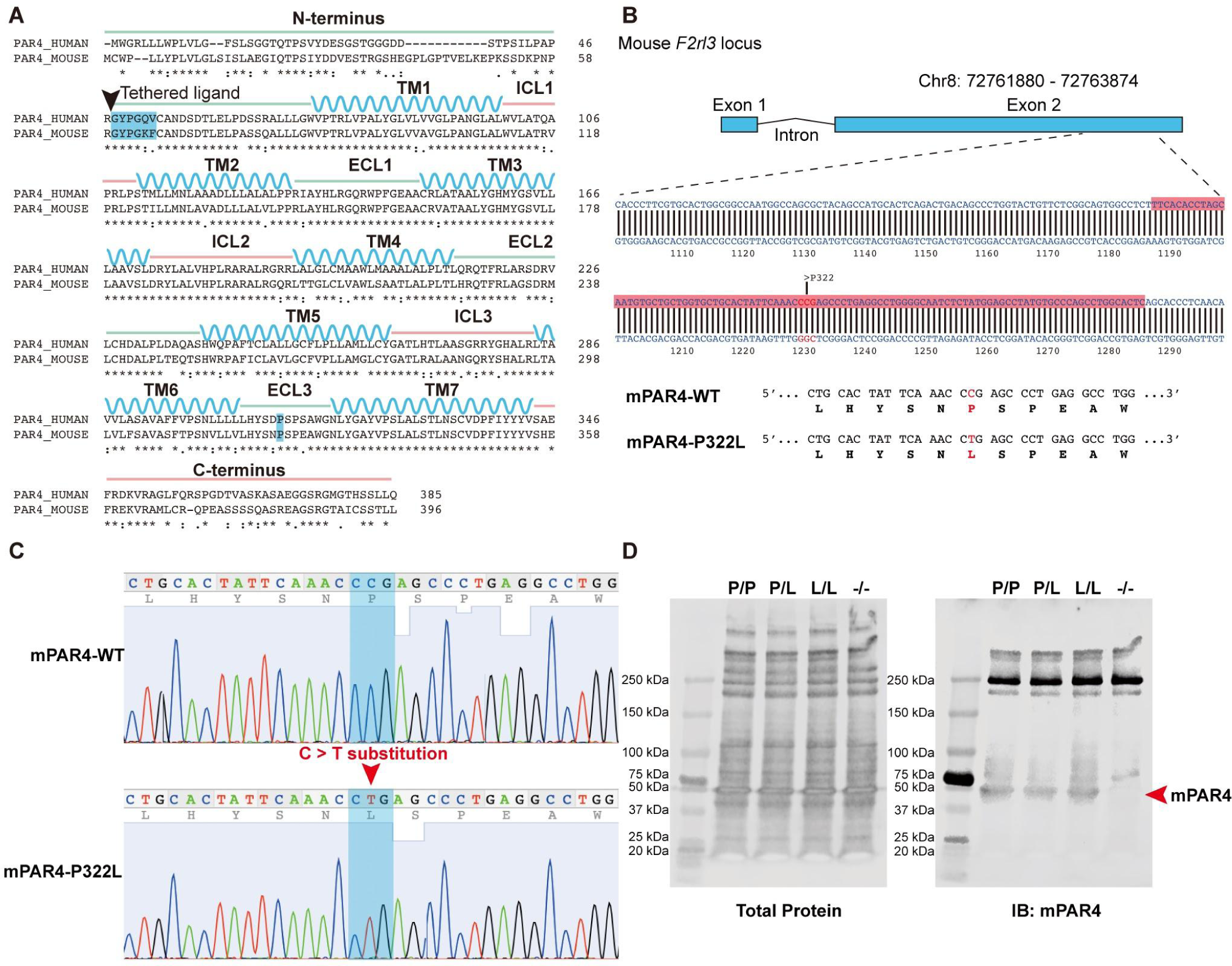
Introduction of the C>T substitution in *F2rl3* locus. **(A)** Sequence alignment of human PAR4 and mouse PAR4. P310 at ECL3 (highlighted in blue) in humans is equal to P322 in mice. **(B)** gRNAs were targeted against the region of the mouse F2rl3 locus highlighted in red, while the red letters indicate the substitution target site. **(C)** Sanger sequencing shows the *F2rl3* locus around the gRNA target site of a wildtype mouse (top) and a PAR4-P322L homozygous mouse (bottom) in which both alleles contained C>T substitutions. **(D)** Protein levels of PAR4 were compared across genotypes using an antibody specific for the mouse protein.

### PAR4-P322L reduced platelet response to PAR4-activation peptide (PAR4-AP, AYPGKF)

To characterize the platelet function of the PAR4^P/L^ and PAR4^L/L^ mice in response to PAR4-specific stimulation, murine platelet rich plasma (PRP) from PAR4^P/P^, PAR4^P/L^ or PAR4^L/L^ littermates was stimulated with 50 – 1600 μM PAR4-activation peptide (AP), AYPGKF-NH_2_. Integrin activation and α-granule secretion were measured by flow cytometry using antibodies specific for activated αIIbβ3 or P-selectin (**Figure 2A-B**). 400 μM PAR4-AP was sufficient to elicit the maximum activation of the platelets from PAR4^P/P^ mice with wild-type PAR4 on their surface, with an EC50 of 180 μΜ for P-selectin and 144 μΜ αIIbβ3. These data are summarized in **Table 1**.

**Figure 2.**
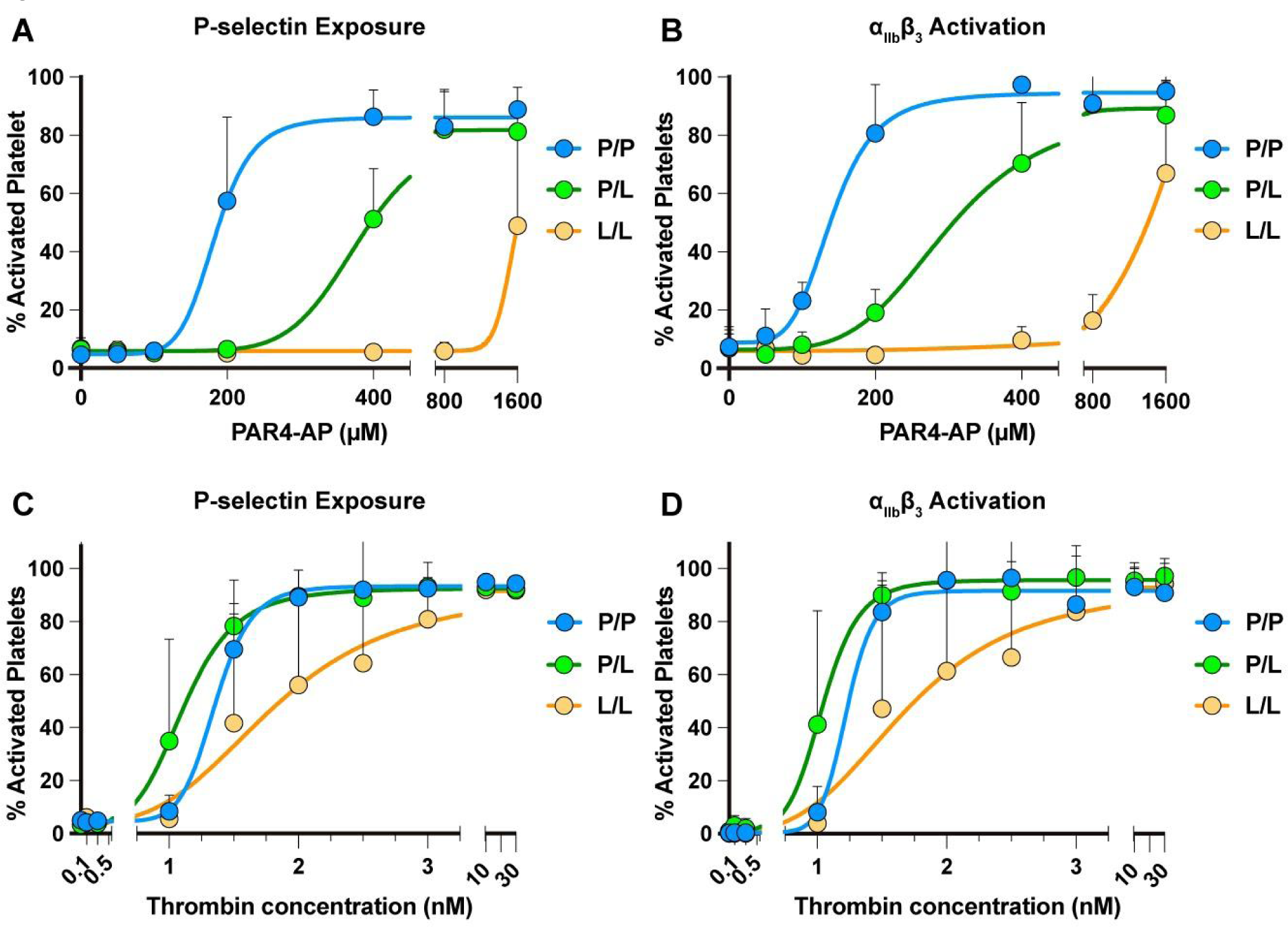
Platelets from PAR4-P322L mice were less responsive to PAR4 agonists. Platelet rich plasma (PRP) from PAR4^P/P^, PAR4^P/L^, PAR4^L/L^ littermates were stimulated with 50-1600 μM PAR4-AP, AYPGKF-NH_2_. (**A, B**). Gel-filtered platelets from PAR4^P/P^, PAR4^P/L^, PAR4^L/L^ littermates were stimulated with 0.1 - 30 nM thrombin (**C, D**). The α-granule secretion (**A, C**) and integrin αIIbβ3 activation (**B, D**) were measured by flow cytometry using antibodies specific for P-Selectin and activated αΙΙbβ3. Data are means ± SD from 5 independent experiments at each concentration for A and B; Data are means from 3 independent experiments at each concentration for C and D.

**Table:** Platelet activation determined by P-selectin positive or integrin αIIbβ3 activation using flow cytometry.

| Table: Platelet activation determined by P-selectin positive or integrin αIIbβ3 activation using flow cytometry. | | | | |
| --- | --- | --- | --- | --- |
| | PAR4-AP (AYPGKF)(μM) | | Thrombin (nM) | |
| | αIIbβ3 activation | P-Selectin | αIIbβ3 activation | P-Selectin |
| PAR4-P/P | 141 (125-160) | 184 (156 - nd) | 1.2 | 1.4 (1.3 - nd) |
| PAR4-P/L | 300 (267-335) | 380 (347 - nd) | 1.0 (0.95-1.2) | 1.1 (1.0-1.3) |
| PAR4-L/L | ~ 8900 | ~ 1544 | 1.7 (1.3 - 2.1) | 1.8 (1.4 - 2.2) |
| Data are presented as EC50 with a ± 95% confidence interval. Undefined confidence intervals are indicated as nd. The maximum response of PAR4-L/L to AYPGKF is greater than 1600 μM; EC50 values are estimated. | | | | |

PAR4^P/L^ mice required 800 μM PAR4-AP to reach maximum activation, with an EC50 of 368 μΜ for P-selectin and 284 μΜ αIIbβ3. There was no difference in the maximum response to the PAR4-AP between the platelets from PAR4^P/L^ and PAR4^P/P^ mice. Further, the platelets from the homozygous PAR4^L/L^ mice showed a pronounced reduction in their response to PAR4 stimulation. When the platelets were treated with the highest dose of PAR4-AP, 1600 μM, we saw only 50% of maximal response with P-selectin exposure and 60% with activated αIIbβ3. Taken together, PAR4-P322L decreased receptor reactivity on mouse platelets, which reduced platelet responsiveness to PAR4 activation peptide.

### PAR4-P322L altered platelet responsiveness to thrombin

Thrombin is the major protease of the coagulation cascade and activates mouse platelets by cleaving PAR4. Gel-filtered platelets from PAR4^P/P^, PAR4^P/L^, or PAR4^L/L^ littermates were stimulated with 0.1 – 30 nM thrombin to determine the response to the endogenous activator. Integrin αIIbβ3 activation and α-granule secretion were measured by flow cytometry using antibodies specific for activated αIIbβ3 or P-selectin, respectively (**Figure 2C-D**). Platelets from PAR4^P/P^ and PAR4^P/L^ mice reached maximum activation at 2 nM thrombin, with an EC50 of 1.1 nM for P-selectin and 1.0 nM for αIIbβ3. Platelets from PAR4^L/L^ mice required more thrombin to reach maximum activation, with an EC50 of 1.8 nM as measured by P-selectin, and 1.7 nM for αIIbβ3. These findings indicated that PAR4-P322L significantly reduced platelet response to thrombin.

### PAR4-P322L did not affect platelet responsiveness to ADP and convulxin

We then determined whether PAR4-P322L altered other signaling pathways by testing platelet response to non-PAR4 agonists. Diluted murine PRP from PAR4^P/P^, PAR4^P/L^ or PAR4^L/L^ littermates were stimulated with ADP (2.5 – 20 μM), the agonist for P2Y1 and P2Y12 receptors, (**Figure 3A-B**) or convulxin (5 nM and 20 nM), a ligand for the platelet collagen receptor, glycoprotein VI. (**Figure 3C-D**) Platelet activation was characterized by integrin αIIbβ3 activation and α-granule secretion measured by flow cytometry. The platelets from the PAR4^P/L^ or PAR4^L/L^ mice had the same response to ADP and convulxin as wild-type mice (PAR4^P/P^). Therefore, PAR4-P322L did not change platelet responsiveness to ADP or convulxin, indicating that PAR4-P322L impaired PAR4-mediated platelet reactivity without impacting other signaling pathways.

**Figure 3.**
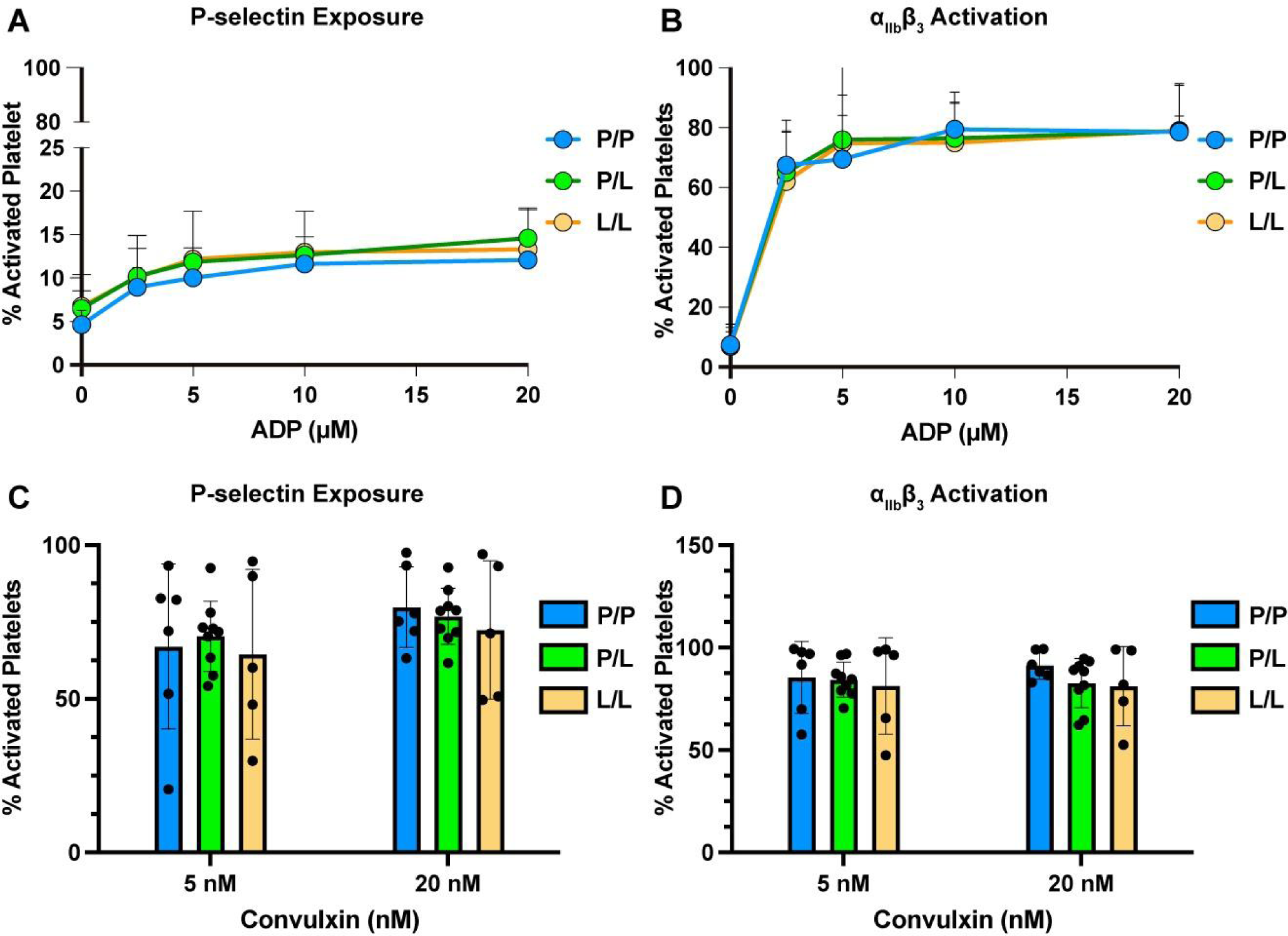
Reactivity to non-PAR4 agonists was unaffected in platelets from PAR4-P322L mice. Platelet rich plasma (PRP) from PAR4^P/P^, PAR4^P/L^ and PAR4^L/L^ littermates were stimulated with 2.5 - 20 μM ADP (**A, B**), or 5 nM or 20 nM convulxin (**C, D**). The α-granule secretion (**A, C**) and integrin αIIbβ3 activation (**B, D**) were measured by flow cytometry using P-selectin and activated αIIbβ3 specific antibodies. Data are means ± SD from 5 independent experiments at each concentration. Dots represent individual mice.

### PAR4-P322L diminished thrombin-mediated platelet aggregation

Since PAR4-P322L reduced platelet integrin activation and granule release in response to thrombin, we next evaluated how this mutation impacted platelet aggregation. Wild-type (PAR4^P/P^) platelets reached a maximum aggregation of 74% (± 6%) response with 1 nM of thrombin while PAR4^P/L^ platelets required 3 nM to reach the same degree of aggregation (**Figure 4A-B**). PAR4^L/L^ platelets required 3 nM to reach maximum aggregation of only 65% (± 5%), 9% lower than that of wild-types (**Figure 4C-D**). A significant decrease in the area under the curve indicated that overall platelet aggregation was negatively impacted by PAR4-P322L, and the rate of aggregation decreased as PAR4 reactivity did as well (**Figure 4E-F**). Altogether, this showed that the PAR4-P322L mutation significantly impaired platelet aggregation.

**Figure 4.**
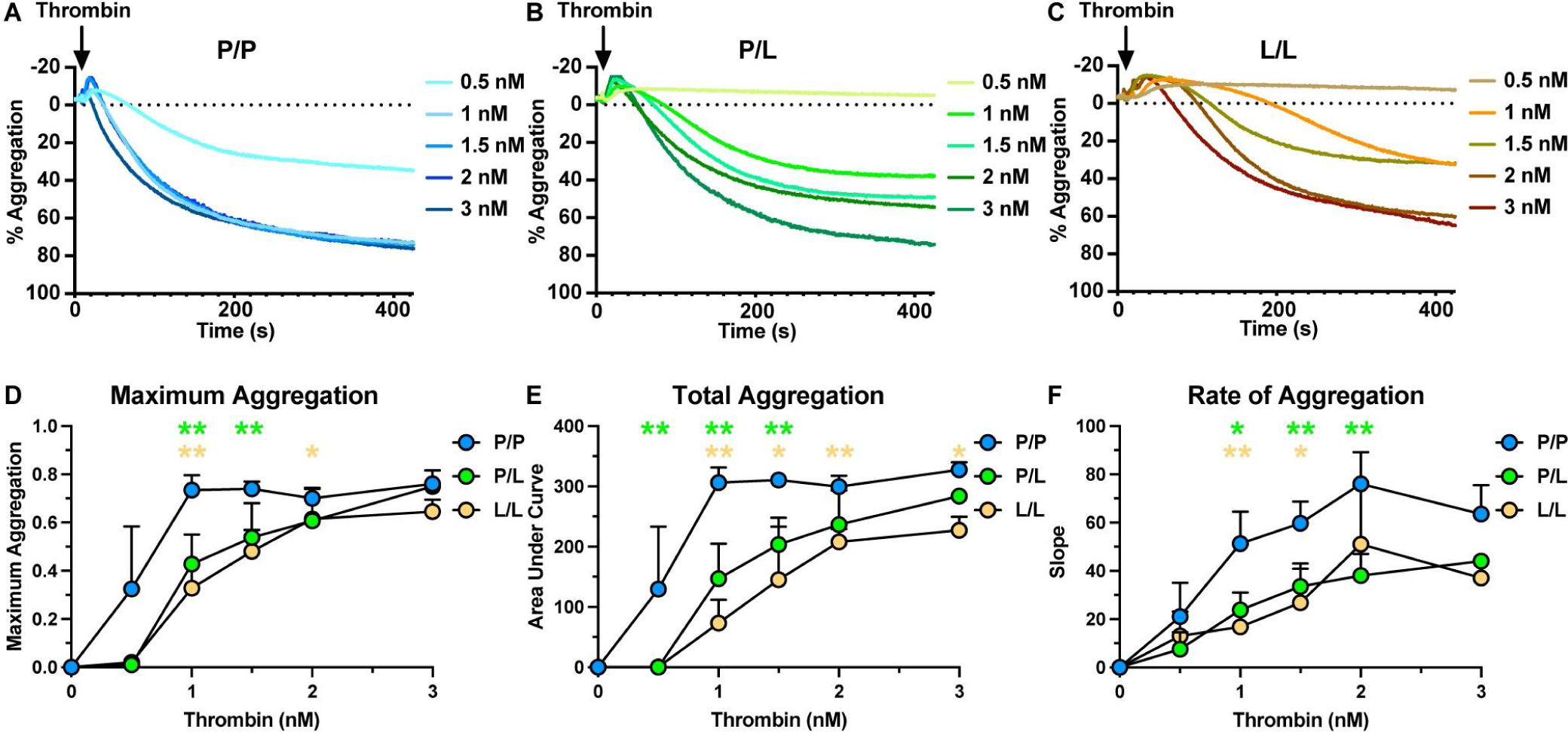
Aggregation of platelets from P322L mice had a reduced response to thrombin. Representative tracing of thrombin-mediated platelet aggregation (**A-C**). Gel-filtered platelets from PAR4^P/P^ (**A**), PAR4^P/L^ (**B**) and PAR4^L/L^ (**C**) littermates were stimulated with 0.5 – 3 nM thrombin. (**D**) Maximum aggregation of gel-filtered platelets from PAR4^P/P^, PAR4^P/L^ and PAR4^L/L^ were compared in response to 0.5 – 3 nM thrombin stimulation. (**E**) Area under curve of PAR4^P/P^, PAR4^P/L^ and PAR4^L/L^ platelet aggregation was compared in response to 0.5 – 3 nM thrombin stimulation. (**F**) The aggregation rate of PAR4^P/P^, PAR4^P/L^ and PAR4^L/L^ platelet aggregation was compared in response to 0.5 – 3 nM thrombin stimulation. Data are representative of 3 independent experiments. Dots represent individual mice.

### PAR4-P322L extended tail bleeding time

The hemostatic plug is formed in response to vascular injury with the goal of preventing blood loss. The tail bleeding assay was used to characterize the hemostatic function of the homozygous PAR4^L/L^ and heterozygous PAR4^P/L^ mice. Both male and female mice at the age of 8 weeks were used. The body weight ranged from 23-26 g in the male mice and from 16-20 g in the female mice. We measured the time to initial cessation of bleeding within 10 minutes of tail snip (**Figure 5A**). The time to initial cessation of bleeding was unchanged in PAR4^P/L^ mice, which averaged 225 ± 96 seconds, when compared to wild-type PAR4^P/P^ littermates, at 189 ± 117 seconds. PAR4^L/L^ mice did take significantly longer to initially stop bleeding (326 ± 152 seconds). To account for rebleeding due to unstable clots, we also measured the total bleeding time over 10 minutes (**Figure 5B**). The total bleeding time was also unchanged between PAR4^P/P^ and PAR4^P/L^ mice, 262±152 versus 235±84 seconds, respectively.

**Figure 5.**
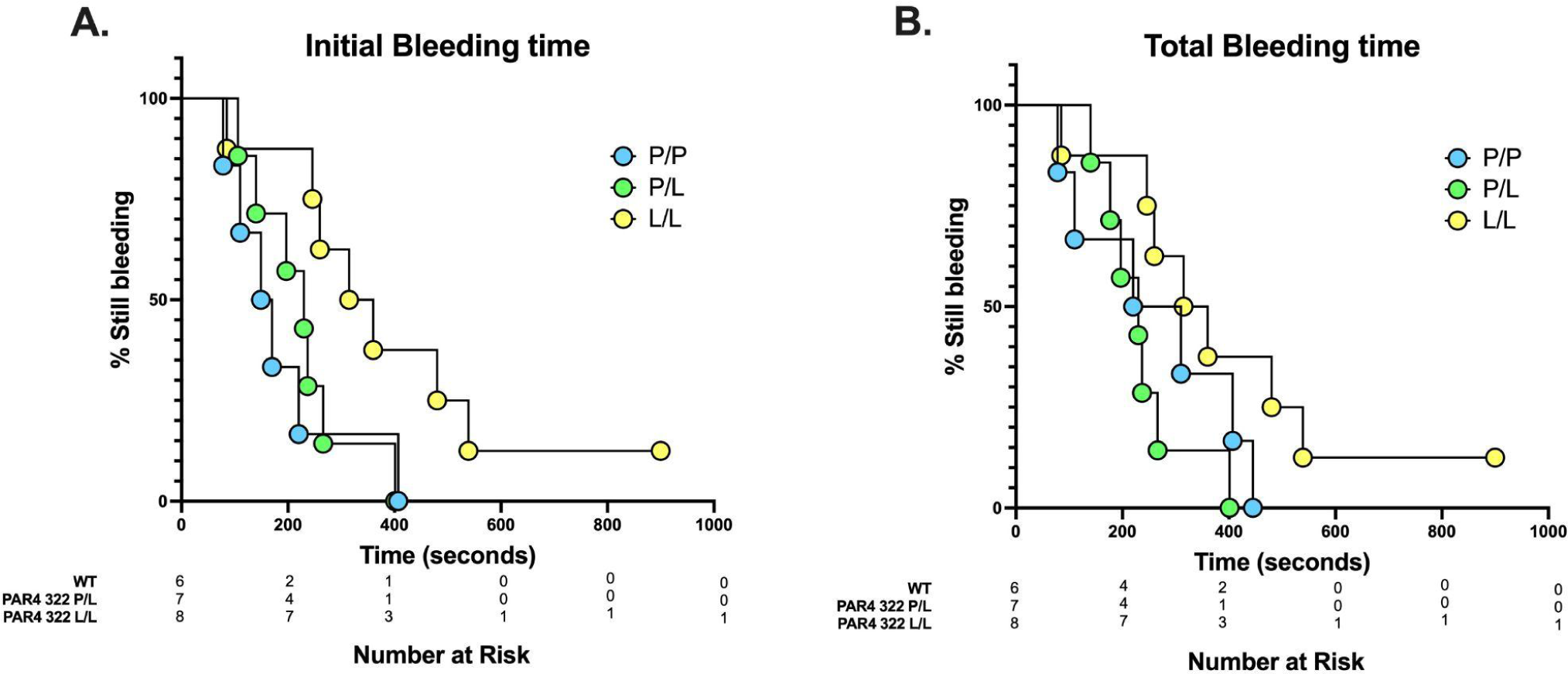
PAR4-P322L mice had extended tail bleeding time. Tail bleeding assay was used to evaluate the impact of the PAR4-P322L on hemostatic function. (**A**) Initial bleeding time was defined as the first time observing the stop of the bleeding regardless of any re-bleeding. (**B**) Total bleeding time was defined as the sum of bleeding times of all bleeding on/off cycles until a stable cessation occurred (no bleeding for 60 s). The experiment was terminated at 10 minutes. The data were presented as the percentage of mice that were still bleeding at a specified time point.

### PAR4-P322L mice had an increased time to arterial occlusion

Global PAR4 knockouts are protected against arterial thrombosis in the FeCl_3_ model of carotid artery occlusion.[27] Our results are consistent with these reports. All PAR4^P/P^ mice fully occluded by 17 minutes (12 min ± 3.4), while PAR4^-/-^ mice were unable to develop thrombi by 30 minutes (**Figure 6A**). PAR4-P322L mice showed an increased time to occlusion compared with wild-types, with 55% of PAR4^P/L^ and 18% of PAR4^L/L^ mice unable to develop stable thrombi. Interestingly, there was a difference in the trends between male and female mice. Male PAR4^P/L^ mice showed similar time to occlusion when compared to wild-types, while 29% of PAR4^L/L^ mice were unable to fully occlude (**Figure 6B**). In females, PAR4^L/L^ mice were similar to wild-types, however 83% of female PAR4^P/L^ mice were unable to develop stable thrombi (**Figure 6C**). Representative images over 30 minutes for each genotype are shown (**Figure 6D**).

**Figure 6.**
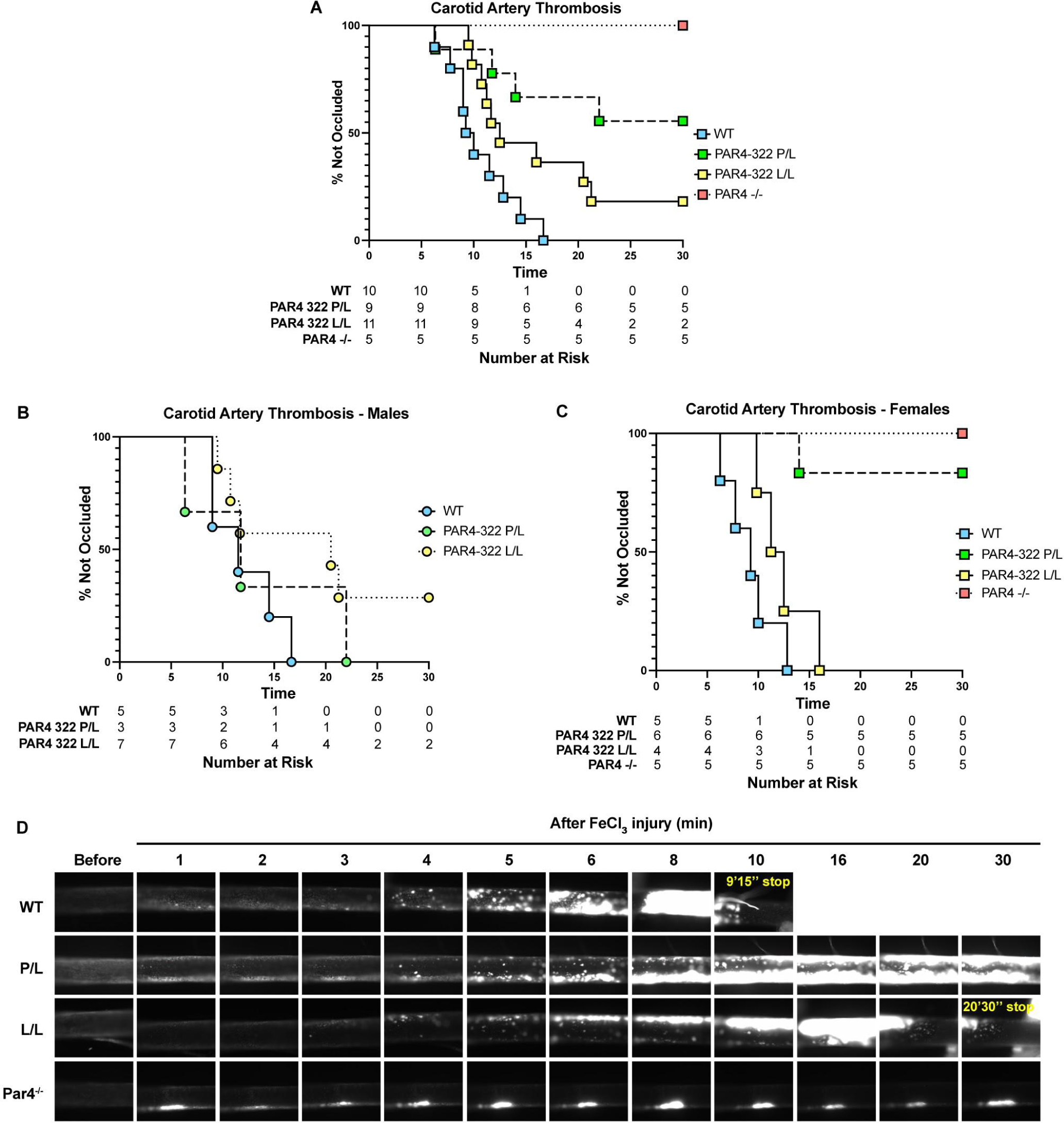
PAR4-P322L mice have longer arterial occlusion times. Arterial thrombosis in our mice was assessed with the ferric chloride-induced carotid artery injury model. (A) The time to occlusion was visually determined as the moment blood flow stopped. The time to complete occlusion was determined in males (B) and females (C). Rhodamine 6G was used to label white blood cells and platelets in real-time over 30 minutes. Representative images are shown. (D).

## Discussion

PAR4 is one of the key thrombin receptors on the surface of human platelets. This G-protein coupled receptor (GPCR) is activated via a unique tethered ligand mechanism and relays extracellular stimulation across the cell membrane[28]. Our previous structural studies demonstrated that a PAR4 single nucleotide polymorphism (SNP) in the extracellular loop 3 (ECL3), PAR4-P310L (rs2227376), has a reduced receptor function in exogenously expressing HEK293 cells[19].

Here, we designed a mouse model to investigate how the hypo-reactive leucine allele at PAR4-P310 impacts PAR4-mediated platelet function. A point mutation was introduced into the mouse PAR4 gene, F2rl3, via CRISPR/Cas9 to create PAR4-P322L, the mouse homolog to human PAR4-P310L. Platelets from the heterozygous (PAR4^P/L^) and homozygous (PAR4^L/L^) mice showed a significantly reduced response to both PAR4-activation peptide (PAR4-AP) and thrombin stimulation compared to wild-type mouse platelets. The PAR4-P322L mutation had no impact on other signaling pathways, such as P2Y12 or GPVI, but did extend tail bleeding time in homozygous PAR4^L/L^ mice. The mutation did show a significant impact on arterial thrombosis and extended clotting time in the FeCl_3_ model of carotid artery injury.

Further study of the PAR4-P310L variant will reveal more about how mutations in GPCRs impact structure and function. PARs belong to the class A GPCRs, which all share an architecture of seven transmembrane domains connected by extracellular and intracellular loops. The original hypothesis for the role of the extracellular loops (ECLs) was that they act as linkers to connect the transmembrane domains together. However, more compelling evidence in recent years indicates that those loops are key mediators for many critical aspects of GPCR function[3]. For PARs, like many other class A GPCRs, ECL2 has been shown to play a key role in ligand recognition and interaction in PAR1, PAR2, and PAR4[7]. Interestingly, our more recent structural approach using amide hydrogen deuterium exchange coupled with mass spectroscopy (HDX-MS) revealed an unrecognized role of ECL3 in PAR4 activation[19]. Based on our studies using PAR4 mutants in cells, we propose the coordinated movement of ECL3 is required for PAR4 activation. The rigidity of the loop is conferred by two conserved prolines (Pro310 and Pro312), and is essential for the dynamic rearrangement of ECL3. We mutated the “rigid” proline at 310 position on ECL3 to a more “flexible” leucine, which led to a PAR4 hypo-reactive to both thrombin and PAR4-AP stimulation, as measured by intracellular calcium mobilization. We hypothesized that the Leu310 blunts PAR4 activation and creates a hypo-reactive receptor, which subsequently decreases platelet responsiveness to PAR4 stimulation.

PAR4 serves as a major thrombin signal initiator on platelets, and polymorphisms other than P310L have been described and implicated in diseases like stroke and thrombosis. The rs773902 SNP (PAR4-120Ala/Thr) results in an amino acid switch at position 120, where Thr120 makes PAR4 more sensitive to agonists than the Ala120 variant.[20] Ala120 is located within transmembrane domain 2, an area close to the ligand binding site.[19] The PAR4-Thr120 variant renders the receptor less sensitive to PAR4 antagonists and worsens stroke outcomes in a humanized mouse model.[21,23] In populations where this SNP is more prevalent, like those of African ancestry (61.5% against 20.6% in people of European ancestry), these individuals are less responsive to antiplatelet agents and are at a higher risk of ischemic stroke.[23] A less common PAR4 variant is the SNP at rs2227346 (PAR4-296Phe/Val). Val296 has only been seen in black individuals, and this variant abolished hyperactivation related to PAR4-Thr120. Phe296 is located within a conserved switch region in transmembrane domain 6 that is important for GPCR activation[20]. Together, this is evidence that SNPs in PAR4 impact PAR4-mediated platelet activation that can be clinically relevant when treating patients. A GWAS meta-analysis done to identify genetic variants associated with venous thromboembolism (VTE) using the INVENT consortium database provided us evidence that rs2227376 is associated with a 15% reduction in relative risk for VTE.[19,29] The minor allele frequency for rs2227376 in the INVENT consortium database was 0.015. With our growing knowledge of platelets and their role in the early stages of VTE, it is likely that PAR4-P310L is directly impacting platelet function in these individuals that results in a degree of protection from thrombosis.

The conserved nature of PAR4 across species, and specifically ECL3, reinforces the idea that this loop is essential for function. It also allowed us to use CRISPR to introduce a point mutation at PAR4-P322 in mice, homologous to human P310, to study this mutation’s impact on physiological functions in more detail. Studying the platelets from these mice has allowed us to see what reduced PAR4 function means for platelet activation in hemostasis and thrombosis. PAR4-mediated platelet activation has been extensively studied in the context of primary hemostasis, which is a complicated process that requires the participation of many macromolecules. Thrombin-mediated PAR4 activation is just one aspect of hemostasis. In fact, based on the results of our platelet function experiments, it was clearly demonstrated that ADP and collagen signaling pathways were unaffected by the PAR4-P322L mutation. This may explain why there was no significant difference in tail bleeding among the PAR4^P/P^, PAR4^P/L^ and PAR4^L/L^ mice while PAR4 signaling was clearly impaired. This further highlights the potential of PAR4 as a target for antithrombotic therapies which aim to inhibit thrombosis without impairing hemostasis.

PAR4 is highly conserved between species, but there are key differences in signaling and function that should be considered when interpreting our findings. PAR4 is present on platelets in both mice and humans, but because it is not an efficient thrombin substrate, it requires a cofactor for low concentrations of thrombin.[30–32] Human platelets express PAR1, while mouse platelets express PAR3.[33] A benefit of studying PAR4 in mice is that it allows the protein to be studied in the absence of PAR1, which is much more sensitive to thrombin. In mice, PAR3 acts as a cofactor for PAR4 activation at low levels of thrombin, and its presence is not required for thrombin-mediated platelet activation.[32,34] Our data in mouse platelets suggests the P322L mutation predominantly impacts thrombin-mediated PAR4 activation at low thrombin concentrations, and we are not seeing compensation for this loss from PAR3. Based on the location of the mutation and ECL3, it is also unlikely that this mutation impacts PAR4’s ability to dimerize with itself, other PARs, or receptors like P2Y12.[35] Recent studies with mice expressing human PAR4 also suggest that human and mouse PAR4 are more different than previously understood.[36] Altogether, they showed that human PAR4 is more responsive to lower concentrations of thrombin and activated platelets to a greater degree. It is likely that the P322L mutation in a more sensitive, humanized PAR4 would more greatly impact its signaling, especially around low concentration of thrombin. With the development and validation of this mouse model, it would be beneficial to study this mutation in a humanized setting.

Using our mouse model permits us to study the impact of reduced PAR4 function in a more complex, physiologically relevant system than cell-based assays allow. Our ex vivo data from platelets with the P322L mutation show a dramatic reduction in their response to PAR4 stimulation. Specifically, our platelet data suggests a gene-dosage response where the decrease in reactivity is more pronounced in those with two copies of the leucine allele (PAR4^L/L^) compared to those with one (PAR4^P/L^). In the FeCl_3_ model of carotid artery thrombosis, however, heterozygous PAR4^P/L^ mice showed a more dramatic phenotype with 55% of mice unable to develop a stable thrombi in comparison to 18% of PAR4^L/L^ mice. Importantly, we saw a distinct difference in clot development between the male and female PAR4^P/L^ mice. 83% of female PAR4^P/L^ mice formed unstable thrombi that were unable to fully occlude the artery, which was a phenotype not seen in the PAR4^P/L^ males. PAR4 has been extensively characterized in platelet function and hemostasis, and there is no recorded difference in PAR4 function between males and females. If there is an impact of sex on PAR4 function, it is a subtle phenotype seen only when PAR4 function is reduced, as we model in our P322L mice. Another possibility is that there is another cell type at work in our model that is impacted differently by the mutation. Importantly, the PAR4-P322L mutation in our mouse model is global, which means it is present across cell types where PAR4 is naturally expressed, like endothelial cells. Once the impact of PAR4-P322L on platelet function is fully understood, we can extend our focus to its effect on other cell types.

We know from studies with systemic PAR4 knockout (PAR4^-/-^) mice that thrombin-mediated PAR4 activation drives platelet aggregation and granule release, ultimately driving the formation of the hemostatic plug.[37] PAR4^-/-^ mice also exhibit impaired clot development in several models of arterial thrombosis and show a severe reduction in platelet activation and recruitment to the growing thrombus.[27] Recently, researchers have been able to develop a mouse model with PAR4 deletion specifically on platelets[38]. These mice, similar to the systemic knockouts and our P322L mutants, showed an increased time to occlusion in the FeCl_3_ carotid artery injury model. There was also a negative impact of platelet-specific PAR4 deletion on venous thrombosis. We know that individuals with the PAR4-P310L SNP are at a reduced risk of VTE, and growing evidence shows that platelets help drive clot development.[39–41] In fact, platelet dysfunction and platelet-neutrophil aggregation have been found to be possible risk factors for VTE.[42,43] Knowing that PAR4 activation promotes platelet-neutrophil binding and thrombin generation increases in VTE, it is likely that thrombin-mediated PAR4 activation is driving thrombosis via platelets.[44,45] Going forward, we can use our PAR4-P322L model to further study the underlying mechanism of PAR4 contributions to VTE alone and in combination with other pathologies to highlight the benefit of targeting PAR4 therapeutically.

## Acknowledgements

Funding for this study was provided by the National Institutes of Health (NIH) National Heart Lung and Blood Institute (NHLBI) (HL098217 and HL154026 M.N.), the Ruth L. Kirschstein Predoctoral Individual National Research Service Award (F31HL162548), and the American Heart Association (AHA) Predoctoral (18PRE33960396 X.H.) and Postdoctoral Fellowships (897185 X.H.). W.L. is supported by R15HL145573. Funding for flow cytometer is supported by Center for AIDS Research (CFAR) (NIH P30AI036219).

## Author Contribution

Study Design: XH, EK, WL, MN Data Collection: XH, EK, MF, WL, WJ Data Analysis: XH, EK, MF, WL, MN Drafting Manuscript: XH, EK, MN Critical Revisions: SR, SM, RC, DL

